# Structure-function analysis of lithium-ion selectivity of voltage-gated sodium channel

**DOI:** 10.1101/2025.05.26.656081

**Authors:** Yuki K. Maeda, Kentaro Kojima, Tomoe Y. Nakamura, Toru Nakatsu, Katsumasa Irie

## Abstract

Voltage-gated sodium channels (Navs) selectively conduct Na^+^ to generate action potentials. Na^+^ permeates Navs with significantly higher efficiency than many other cations, but Li^+^ can also permeate Navs to a comparable extent as Na^+^. It had been known that Li^+^ in blood enters cell via Navs and effects beneficially on various neuropathies. However, the molecular basis of the high Li^+^ selectivity of Navs had been unclear. In this study, using a prokaryotic Nav, we successfully created the first Nav mutant that is more selective for Li^+^ than for Na^+^. Electrophysiological and crystallographic analyses revealed the critical determinants of high Li^+^ selectivity: the strong electrostatic interaction between the ion pathway and hydrated ions, and the smaller number of hydration water exchanges within the ion pathway. Additionally, the extensive interactions around the ion pathway were shown to support monovalent cation selectivity. New drug directions based on the molecular basis for Li^+^ permeation may target various neurological disorders and clarify the broader biological effects of lithium.

## Introduction

Voltage-gated sodium channels (Navs) are membrane proteins that open in response to depolarization of the membrane potential (Hille, 2001). Navs conduct Na^+^ ions inward quickly (∼10^7^ ions/sec) to initiate and propagate rapid action potentials (Catterall, 2023; Hodgkin & Huxley, 1952). The efficiency of Na^+^ permeation of Navs is significantly higher than that of many other cations (Na^+^ ≈ Li^+^ >>K^+^, Cs^+^, Ca^2+^), so Navs enable the Na^+^ selective translocation across cell membranes in response to stimuli (Campbell, 1976; Hille, 1972; Pappone, 1980). This property makes Navs key players in nerve conduction, muscle contraction, secretion, neurotransmission, and many other processes.

The high Na^+^ selectivity of Navs is well known, but Navs also conduct lithium ion (Li^+^) with similar efficiency as with Na^+^ (Hille, 2001). Although its physiological role is rarely understood, lithium preparations are in use for the treatment of manic, hypomanic, and depressive episodes with bipolar disorder (Shorter, 2009; Hart, 2024; Cade, 1949; Carvalho *et al*, 2015; Fountoulakis *et al*, 2016; Malhi *et al*, 2015; Nestsiarovich *et al*, 2022; Verdolini *et al*, 2021). Further beneficial actions of lithium have also been revealed, including neuroprotectivity in Alzheimer’s disease, reduction of mitochondrial dysfunction, and modulation of apoptosis autophagy (Almeida *et al*, 2022; Bortolozzi *et al*, 2024; Chiu *et al*, 2013; Guttuso *et al*, 2019; Haupt *et al*, 2021; Lauterbach; Lazzara & Kim, 2015; Leeds *et al*, 2014; Scheuing *et al*, 2014; Shim *et al*, 2023; Singulani *et al*, 2024; Vallée *et al*, 2021; Vo *et al*, 2015). Li^+^ in the blood, liberated from lithium preparations, is assumed to enter cells via Navs (Timmer and Sands, 1999) and certainly affects various cellular systems (Hart, 2024; Schrauzer, 2002). While the beneficial effects of lithium are well understood, concerns have also been raised about bioaccumulation toxicity, according to the increase of environmental lithium from industrial products, as represented by Li^+^ batteries (Chevalier *et al*, 2024). In vitro, harmful effects of lithium have been reported, such as neurological, immune, and cardiovascular disorders, as well as fetal diseases (Chevalier *et al*, 2024). However, there are few reports on the mechanism by which Li^+^ enters the cell, i.e., the molecular basis by which Li^+^ permeates Navs.

Na^+^ and Li^+^ have different ionic properties, so it is uncertain how they achieve comparable ion permeability in Navs. In mammalian Navs, the pore-forming subunit (α_1_) consists of four repeats, which are formed by six-fold transmembrane segments (S1-S6) in each repeat (Abriel & Kass, 2005; Catterall, 2000) (Figure 1A). Each repeat consists of two domains: S1-S4 detects membrane voltage as the voltage-sensing domain, and S5-S6 forms the ion pore by facing each other, thereby building a pore domain. The ion pore consists of an outer funnel-like vestibule, a selectivity filter (SF), a central cavity, and an intracellular activation gate. In particular, the SF is especially protruded into the ion pore by the support of the P1 and P2 helices that flank the SF in the pore-loop between S5 and S6. The permeation efficiency of each ion is mainly determined by the five amino acid residues of the SF, which narrow the ion pore the most (Hille, 2001). However, the asymmetric and tortuous configuration of vertebrate Navs makes it challenging to fully understand the extent of ion permeation processes.

**Figure 1.**
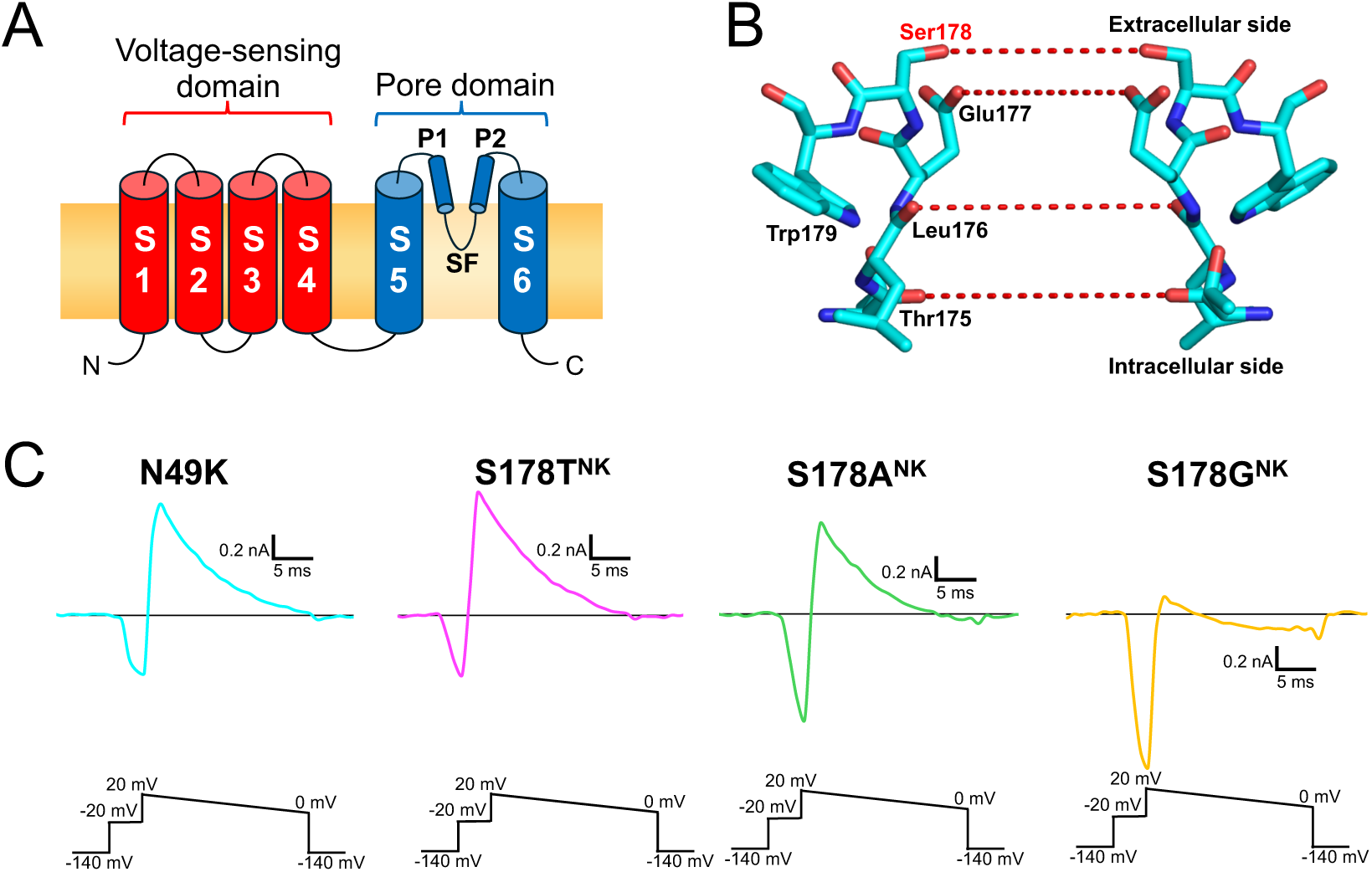
The selectivity filter of NavAb and representative current traces of NavAb N49K, S178T^NK^, S178A^NK^ and S178G^NK^ mutants **A)** Schematic diagram of BacNav subunit. Mammalian Navs comprise four homologous repeats. **B)** Side view of the residues 175-179^th^ constituting the selectivity filter of NavAb N49K mutant (PDB code: 8H9W). The red dashed lines represent the hydration water exchange sites that ions pass through, formed by the oxygen atoms of the main and side chains of the SF. **C)** Recordings of the current responses of NavAb N49K, S178T^NK^, S178A^NK^, and S178G^NK^ mutants obtained using the ramp protocol with Li^+^ extracellular solution. Currents were generated by the step pulse of -20 mV from -140 mV holding potential, followed by the ramp pulse with different voltage values (shown at the bottom).

As a substitute for vertebrate Navs, the Na^+^ permeation mechanism of Navs is well explained by the functional analyses and three-dimensional structures of several bacterial Navs (BacNavs) (Ahern *et al*, 2016; Corry & Thomas, 2012; Payandeh *et al*, 2011). BacNavs comprise homotetramers of subunits with a composition that resembles a repeat of vertebrate Navs. Based on phylogenetic analysis, the lineage of BacNavs likely diverged from a progenitor of eukaryotic Navs and voltage-gated calcium channels (Cavs), and the structural features are conserved with each other (Catterall & Zheng, 2015; Liebeskind *et al*, 2013; Zakon, 2012). Importantly, BacNavs also conduct Li^+^ with high efficiency as well as eukaryotic Navs (the permeation ratio of Li^+^ over Na^+^ is 0.6∼0.8 in BacNavs and is 0.9∼1.1 in eukaryotic Navs) (DeCaen *et al*, 2014; Finol-Urdaneta *et al*, 2014; Naylor *et al*, 2016; Hille, 2001). We, therefore, thought that if BacNav mutants with enhanced Li^+^ selectivity were created, it might be possible to identify the critical factors of the permeation of Li^+^ common to eukaryotic Navs.

The chemical relations between the strength of the electrostatic force (which is called “field strength”) and the selectivity of monovalent cations have been well-established experimentally as the Eisenman sequence, which was exhibited by using ion-selective glasses (Hille, 2001; Eisenman, 1962). The Eisenman sequence includes a Li^+^-selective order, but no ion channel had been found that exhibits Li^+^-selectivity. Here, we successfully created the first Li^+^-selective BacNav mutant. Furthermore, electrophysiology and crystallography revealed two key factors contributing to high Li^+^ selectivity: the stronger electrostatic force that attracts hydrated ions to the surface of the SF and the smaller number of hydration water exchanges that occur during passage through the ion pathway. Furthermore, we showed the dense network around the SF supporting monovalent cation selectivity. This is the first study to create a Li^+^-selective channel that matches the most Li^+^-selective order in the Eisenman sequence and to reveal the structural basis of Li^+^ permeation via Navs. This insight into the mechanisms by which Li^+^ enters cells will lead to improved treatments for various neurological disorders and a greater understanding of the biological effects of lithium.

## Results

### Electrophysiological evaluation of ion selectivity

The SF of NavAb consists of “TLESW”, in which Ser178 forms the extracellular entrance of the SF (Figure 1B) (Payandeh *et al*, 2011). When Na^+^ accesses the SF from an extracellular bulk solution, the hydroxyl group of the side chain of the Ser178 coordinates with Na^+^ as the first interaction site in the SF, along with the side chain of the Glu177 and four water molecules (Corry & Thomas, 2012). We first examined the Li^+^ selectivity of the NavAb Ser178 mutants (named S178T^NK^, S178V^NK^, S178A^NK^, and S178G^NK^) in the NavAb N49K mutant (named N49K). The N49K mutation was introduced into all constructs in this study to provide a stable current (Gamal El-Din *et al*, 2013). Three mutants, S178T^NK^, S178A^NK^, and S178G^NK^, showed current responses in the insect-cell expression system (Figure 1C), and S178V^NK^ showed no current. We, therefore, thought it was impossible to verify larger-side-chain mutants than valine. In the Li^+^ extracellular solution, the reversal potentials of S178T^NK^, S178A^NK^, and S178G^NK^ mutants differed from that of the N49K, indicating that Li^+^ selectivity changed by these mutations.

For a correct evaluation of ion selectivity, it is necessary to measure the reversal potential accurately. However, it was difficult to measure the reversal potentials of these mutants, whose currents were immediately inactivated. The T206A mutation is known to slow NavAb inactivation and provide long-lasting currents (Gamal El-Din *et al*, 2019). Therefore, we introduced the T206A mutation to N49K, S178T^NK^, S178A^NK^ and S178G^NK^ mutants (named N49K/T206A, S178T^NK/TA^, S178A^NK/TA^, and S178G^NK/TA^). These NavAb T206A mutants showed slower inactivation and provided more prolonged currents (Figure 2A). Next, the permeation ratio of Li^+^ and Na^+^ (*P*_Li_/*P*_Na_) of N49K/T206A, S178T^NK/TA^, S178A^NK/TA^, and S178G^NK/TA^ mutants was evaluated using the reversal potential in a semi-bi-ionic environment. The reversal potential of S178G^NK/TA^ was highest among the other Ser178 mutants, indicating S178G^NK/TA^ had the highest Li^+^ selectivity (Figure 2B, C). To a lesser extent than S178G^NK/TA^, S178T^NK/TA,^ and S178A^NK/TA^ showed comparable positive shifts of reversal potentials relative to N49K/T206A (Figure 2B, C). *P*_Li_/*P*_Na_ was 2.18 ± 0.14 in S178G^NK/TA^ compared to 0.71 ± 0.02 in the N49K/T206A, 0.96 ± 0.01 in S178T^NK/TA^ and 1.09 ± 0.05 in S178A^NK/TA^ (Table 1). *P*_Li_/*P*_Na_ of S178G^NK/TA^ (2.18 ± 0.14) indicates that the glycine mutant allows Li^+^ to permeate about twice as efficiently as Na^+^, which was approximately triple that of N49K/T206A with Ser178. Next, we also evaluated the selectivity of the other cations (K^+^, Cs^+^, Ca^2+^). *P*_K_/*P*_Na_, *P*_Cs_/*P*_Na_, and *P*_Ca_/*P*_Na_ remained at low values with little change between N49K/T206A and the other three mutants (Figure 3 and Table 1). In other words, there was no increase in non-selective cation permeability, and only Li^+^ selectivity was mainly enhanced in these NavAb Ser178 mutants.

**Figure 2.**
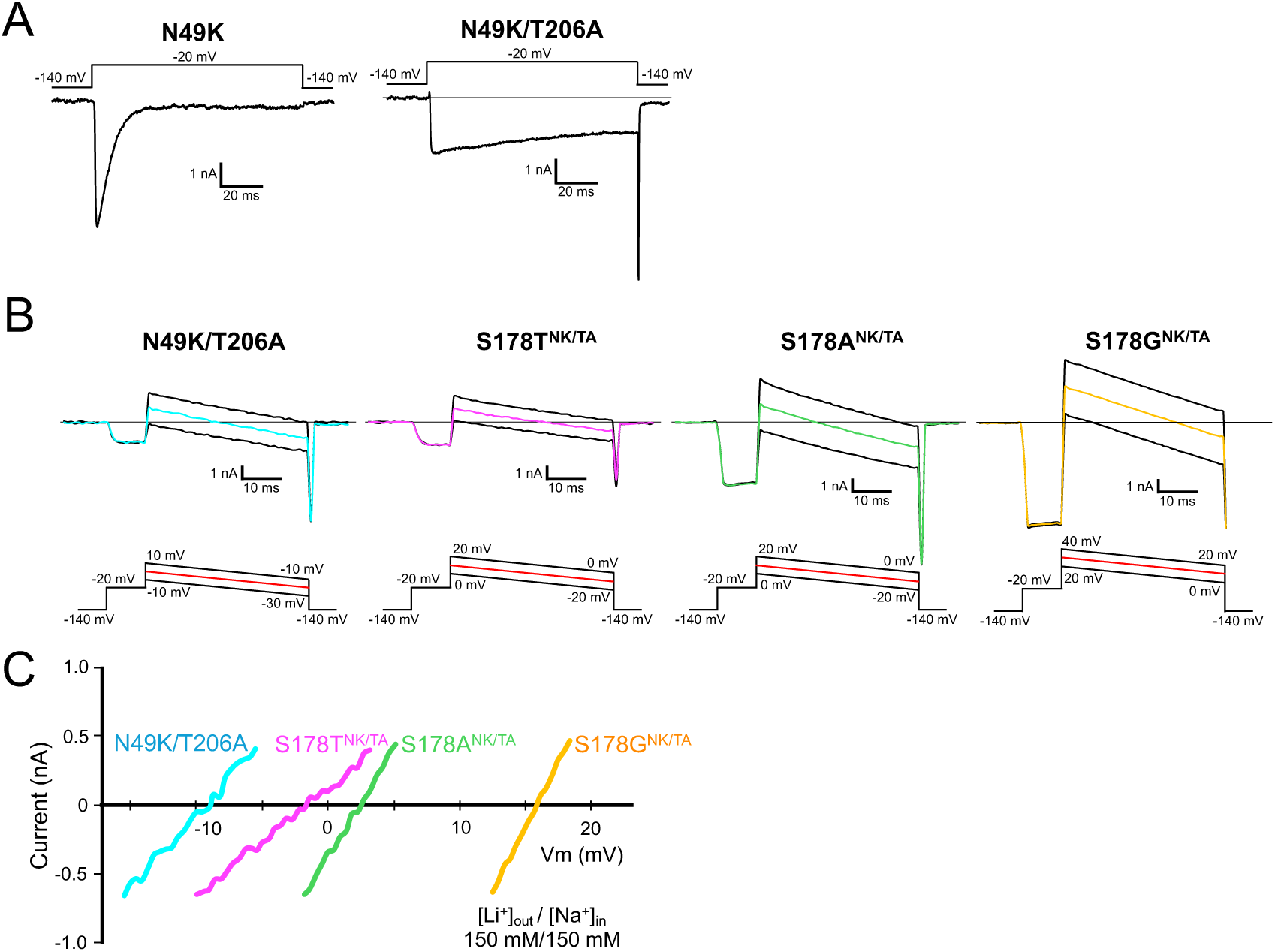
Li^+^ selectivity evaluation in NavAb N49K/T206A, S178T^NK/TA^, S178A^NK/TA^, and S178G^NK/TA^ mutants **A)** Whole-cell currents in N49K when a pulse of -20 mV from -140 mV holding potential was given for 100 ms. **B)** Representative current traces of NavAb N49K/T206A, S178T^NK/TA^, S178A^NK/TA^, and S178G^NK/TA^ mutants were obtained using the ramp protocol. The values of the reversal potential recorded with three different ramp pulses were averaged and used. Currents were measured in 150 mM [Li^+^]_out_ and 150 mM [Na^+^]_in_. **C)** Current-voltage relationship plots of N49K/T206A, S178T^NK/TA^, S178A^NK/TA^, and S178G^NK/TA^ mutants generated by the ramp pulses in 150 mM [Li^+^]_out_ and 150 mM [Na^+^]_in_.

**Figure 3.**
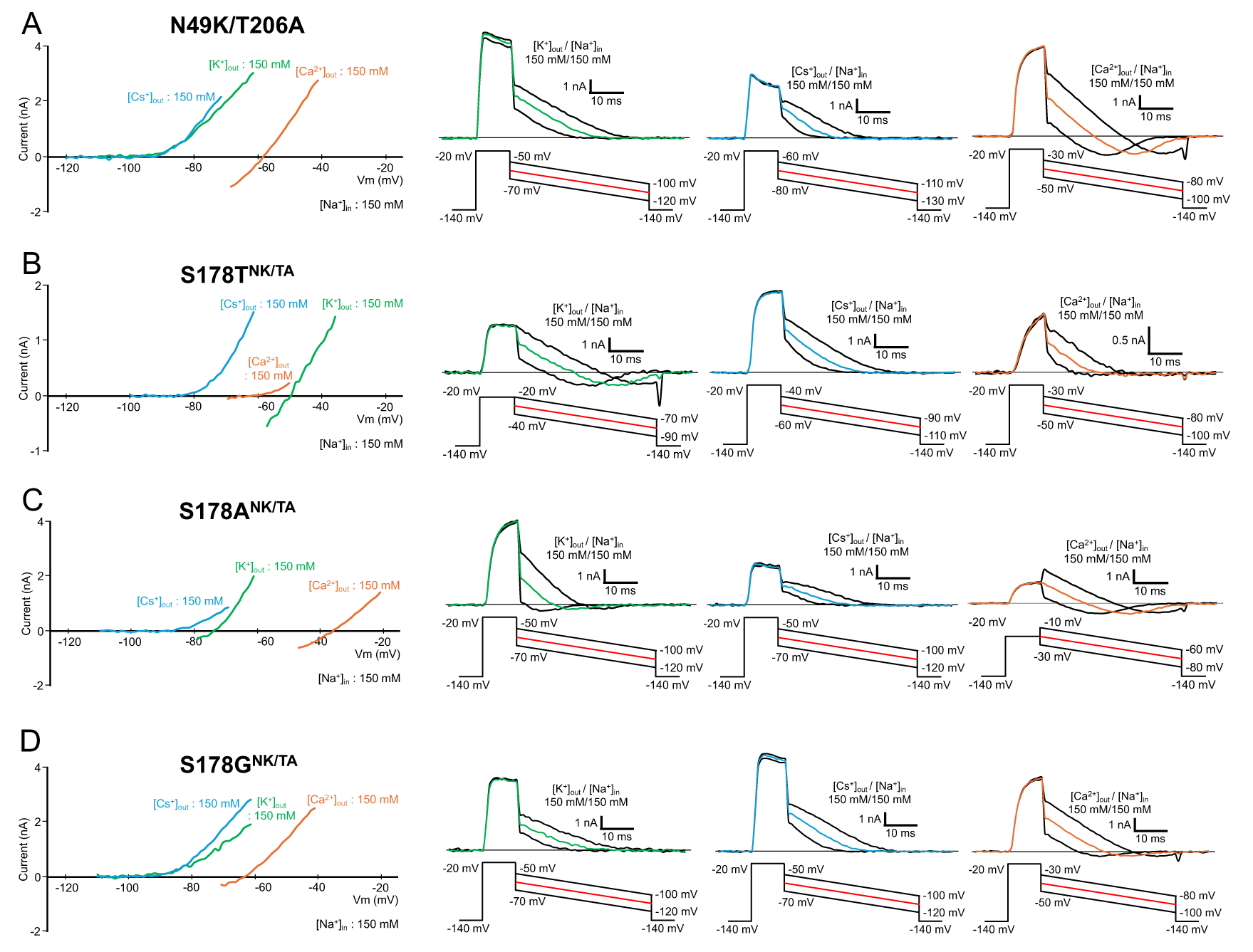
Cation selectivity evaluation in NavAb N49K/T206A, S178T^NK/TA^, S178A^NK/TA^, and S178G^NK/TA^ mutants **A-D)** Current-voltage relationship plots (left) and representative currents (right) used to evaluate the permeability of K^+^, Cs^+^ and Ca^2+^ relative to that of Na^+^ of NavAb N49K/T206A, S178T^NK/TA^, S178A^NK/TA^, and S178G^NK/TA^ mutants. Currents were measured in 150 mM [K^+^]_out_, 150 mM [Cs^+^]_out_ and 100 mM [Ca^2+^]_out_, respectively, and 150 mM [Na^+^]_in_. Currents were generated by the step pulse of -20 mV from -140 mV holding potential, followed by the ramp pulses with different voltage values. The time course of the change of membrane potentials are shown at the bottom of the respective current traces.

**Table 1.** Relative permeability in NavAb N49K/T206A, S178T^NK/TA^, S178A^NK/TA^, and S178G^NK/TA^ mutants. All values are indicated as SEM.

| | $P_{Li}/P_{Na}$ | $P_K/P_{Na}$ | $P_{Cs}/P_{Na}$ | $P_{Ca}/P_{Na}$ |
| --- | --- | --- | --- | --- |
| N49K/T206A | 0.71 ± 0.02<br>(n = 10) | <0.05*<br>(n = 6) | <0.05*<br>(n = 6) | 0.05 ± 0.00<br>(n = 8) |
| S178T<sup>NK/TA</sup> | 0.96 ± 0.01<br>(n = 8) | 0.13 ± 0.01<br>(n = 4) | <0.05*<br>(n = 7) | <0.05*<br>(n = 7) |
| S178A<sup>NK/TA</sup> | 1.09 ± 0.05<br>(n = 11) | 0.05 ± 0.00<br>(n = 8) | <0.05*<br>(n = 6) | 0.12 ± 0.00<br>(n = 6) |
| S178G<sup>NK/TA</sup> | 2.18 ± 0.14<br>(n = 11) | <0.05*<br>(n = 5) | <0.05*<br>(n = 7) | <0.05*<br>(n = 9) |

\*When the reversal potential was below the deactivation potential, $P_M/P_{Na}$ was noted as < 0.05 which is the value calculated from the lowest measurable reversal potential in each construct.

### Crystal structure of mutants

To reveal the molecular mechanisms of the enhancement of Li^+^ selectivity, S178T^NK^, S178A^NK^, and S178G^NK^ mutants of NavAb were crystallized, and these structures were determined at approximately 3.0 Å resolution by the same method as used in our previous study of NavAb (Irie *et al*, 2018, 2023). Crystallization was also performed in the mutants containing T206A, but the resolution of these crystals’ diffraction patterns deteriorated in all T206A mutants. As with the mutants without T206A, the S178T/T206A mutant showed the best resolution in all T206A mutants, but its resolution was only 3.4 Å (Sup. Table 1). The RMSDs of the Cα carbons of S178T^NK/TA^ mutant to S178T^NK^ were 0.23 Å, and Thr206 is located in the channel lumen, distant from Ser178. T206A is considered not to affect the mutation at Ser178, so structural analysis was performed without T206A.

The electron density fitted well to the mutated side chain of the 178^th^ residue of all mutants (Figure 4). Their structures resembled that of the N49K channel (PDB code: 8H9W) (Irie *et al*, 2023). The RMSDs of the Cα carbons of S178T^NK^, S178A^NK^, and S178G^NK^ mutants to the N49K were 0.48, 0.52, and 0.46 Å, respectively. The whole structure of each mutant was, therefore, almost the same as that of the N49K.

**Figure 4.**
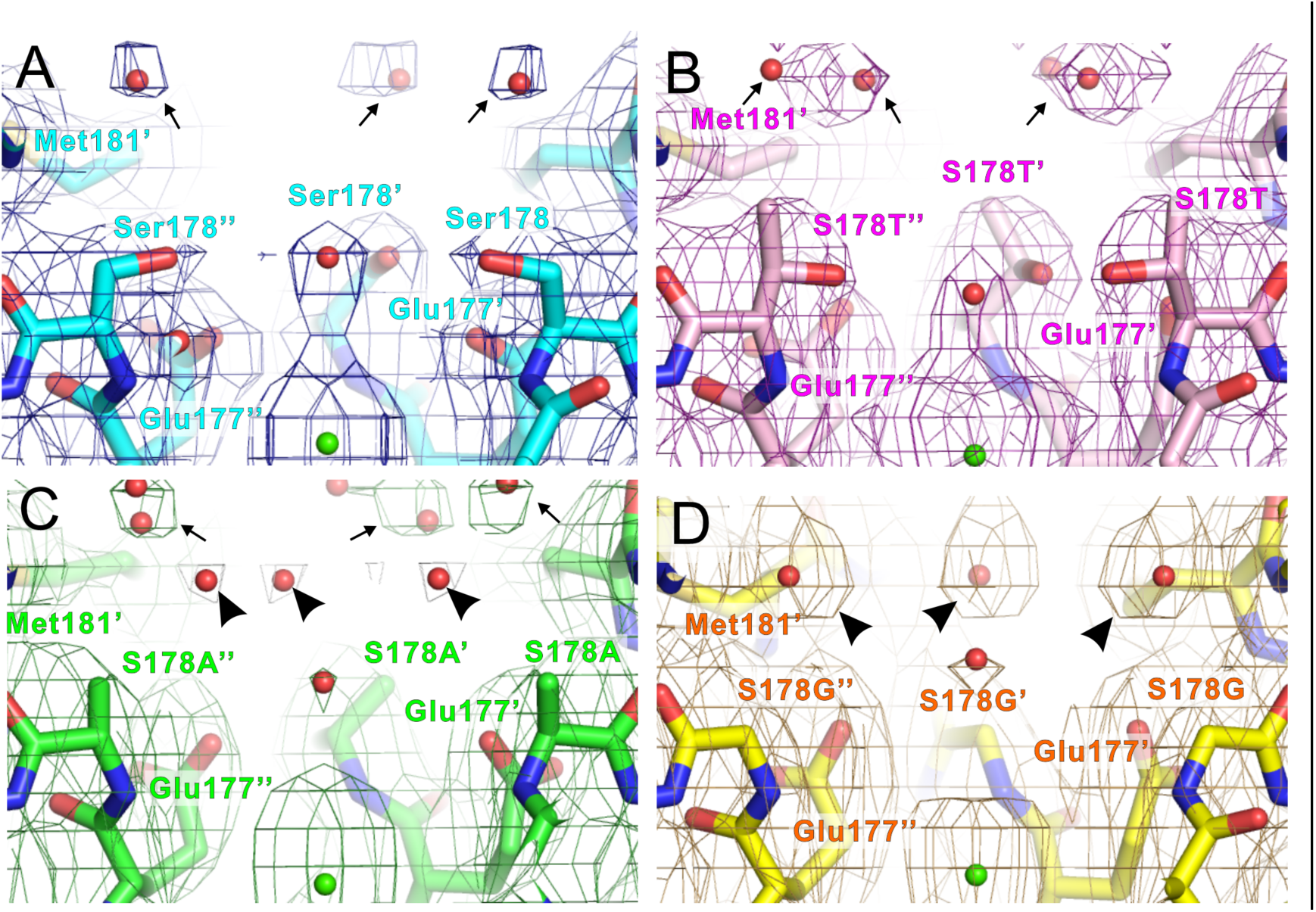
The selectivity filter structure of NavAb N49K, S178T^NK^, S178A^NK^, and S178G^NK^ mutants A-D) Horizontal view of the ion pathway of NavAb N49K, S178T^NK^, S178A^NK^, and S178G^NK^ mutants with the 2*F*_O_ − *F*_C_ electron density map contoured at 1σ, respectively. The carbon atoms of NavAb N49K, S178T^NK^, S178A^NK^, and S178G^NK^ mutants are colored cyan, magenta, green, and yellow, respectively. The electron density of NavAb N49K, S178T^NK^, S178A^NK^, and S178G^NK^ mutants are colored blue, dark magenta, dark green, and orange, respectively. The weak electron density map contoured at 0.5σ are colored gray in NavAb S178A^NK^. The bulk water molecules are indicated by black arrows. The electron density of water molecules are indicated by black arrowheads. The residues of the center subunit are numbered with prime symbols. The Residues of the left subunit are numbered with double prime symbols.

The 178^th^ residue is located at the entrance of the ion pathway. Met181 was located on the bulk solution side of the 178^th^ residue, and water molecules were observed beside this vicinity (Figure 4: arrow). In the S178G^NK^ mutant, the highest Li^+^-selective mutant, the glycine mutation created a large space around the 178^th^ residue position due to the absence of side chains (Figure 4D). The water molecules in the S178G^NK^ mutant were closest to the entrance of the ion pathway among the other mutants (Figure 4D: arrowhead). The S178G mutant exhibited high Li+ selectivity, suggesting that this difference contributes to lithium selectivity. Weak electron density was observed in the S178A^NK^ mutant at a position corresponding to the water molecule near the entrance of the ion pathway in the S178G^NK^ mutant (Figure 4C: arrowhead). Considering that the S178A^NK^ mutant has the second-highest Li^+^ selectivity, water molecules approaching the SF entrance may be favourable for Li^+^ selectivity. When the Ser178 was changed to glycine or alanine, it eliminated the first hydration water exchange site of the SF formed by the hydroxyl groups. At the same time, with one less hydration water exchange site, the first interaction site of the SF then becomes the side chain of Glu177, which is negatively charged. In other words, the enhanced Li^+^ selectivity in the S178G^NK^ and the S178A^NK^ may be related to the smaller number of hydration water exchange sites and the proximity of hydrated ions to the negatively charged sites of the SF.

### The shape of the ion pathway

As the size of the mutated side chain increased, the radius of the ion pore decreased (Figure 5). In the S178T^NK^ mutants, the mutated side chain was the narrowest region, even within the SF, with a length of less than 2 Å (Figure 5B and E). The wild-type Ser residue forms a similarly narrow region, but the radius of this ion pore is more than 2 Å. In the S178A^NK^ and S178G^NK^ mutants, the shape of the ion pore in this region is more expansive because of the smaller-side-chain mutation (Figure 5C and D). These two mutants exhibit improved Li^+^ selectivity, suggesting that the Li^+^ selectivity has been enhanced by widening the pore. On the other hand, this is inconsistent with the fact that S178T^NK^, which has narrower pores, shows a slight improvement in Li^+^ selectivity.

**Figure 5.**
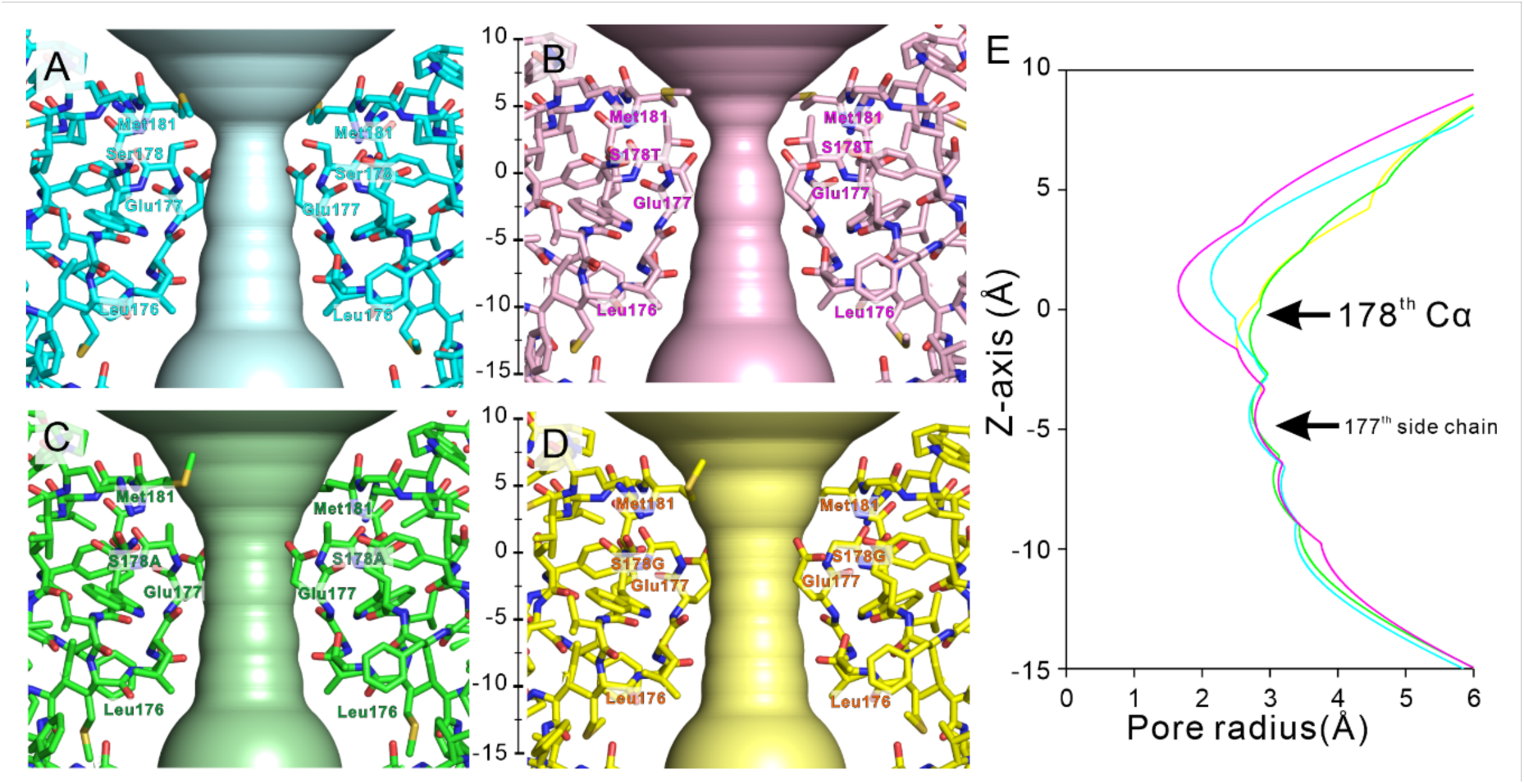
The shapes and radius of the ion pathway of NavAb N49K, S178T^NK^, S178A^NK^, and S178G^NK^ mutants A-D) Horizontal view of the ion pathway of NavAb N49K, S178T^NK^, S178A^NK^, and S178G^NK^ mutants depicted by program hole. The carbon atoms of NavAb N49K, S178T^NK^, S178A^NK^, and S178G^NK^ mutants are colored cyan, magenta, green, and yellow, respectively. E) The pore radius of NavAb N49K, S178T^NK^, S178A^NK^, and S178G^NK^ mutants. The baseline of the z-axis is set to the z-coordinate of the main-chain carbonyl oxygen atoms of the 178^th^ residue. The extracellular direction is plotted as plus, and the cytosolic direction as minus.

Next, we focused on the extensive residues forming ion pores (Figure 6). In the NavAb SF, three residues of the “176-LES-178” form the surface of the ion pathway (Figure 5). The Met181 at the P2 helix was located on the extracellular side of the 178^th^ amino acid residue. In the S178G^NK^ case, the Met181 side chain moved toward the Gln172 side chain at the P1-helix of neighbouring subunit (Figure 6D). The Gln172 is located in the backyard of the ion pore, not facing the ion permeation pathway, but its side chain forms a hydrogen bond with the carbonyl group of the Glu177 main chain (Figure 6). The side chain of Glu177, a carboxylate, forms a critical ion recognition site for ion permeation. The side chain of Arg185 forms a hydrogen bond with the oxygen atom of the side chain of Gln172 and fixes the direction of the amido group of Gln172 to stabilize the hydrogen bond between Glu177 main chain and Gln172 amido group. Similar interactions were observed in the S178T^NK^, S178A^NK^, and S178G^NK^ mutants (Figures 6 B-D). The difference in Met181 observed in S178G^NK^ may, therefore, be responsible for improving Li^+^ selectivity.

**Figure 6.**
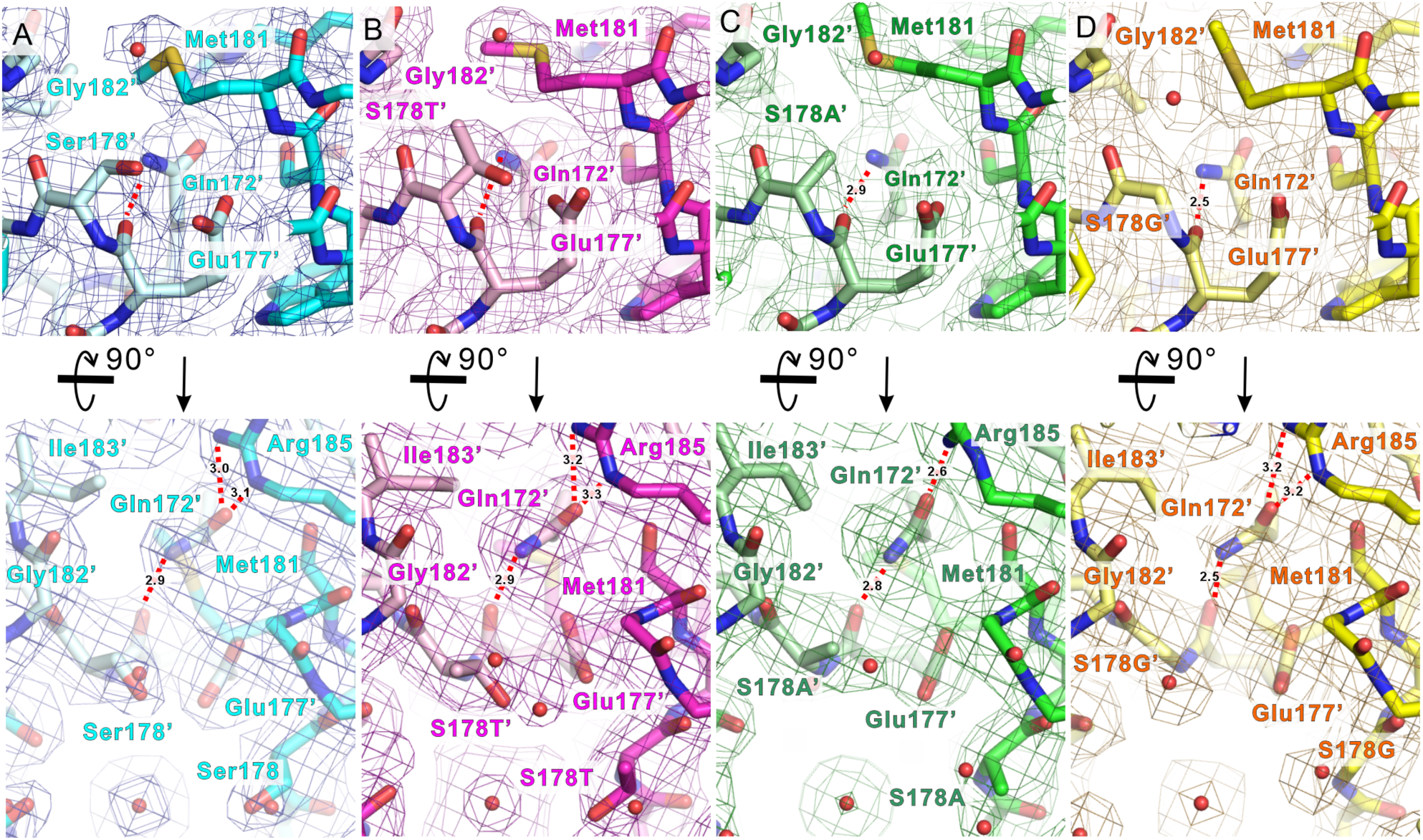
The hydrogen bond network in the backyard of the selectivity filter A-D) Upper: The horizontal view from the center of the ion pathway of the electron densities and protein structures around Met181 and adjacent subunit’s Glu177 and Gln172 of the selectivity filter of NavAb N49K, S178T^NK^, S178A^NK^, and S178G^NK^ mutants, respectively. These figures correspond to the enlarged view of the subunit at the back center in Figure 4. The 2*F*_O_ − *F*_C_ electron density map contoured at 1σ (blue, dark magenta, dark green, and orange). Lower: The vertical view from the extracellular side of the electron densities and protein structures around Met181, indicating Arg185 interaction was lost in S178G^NK^.The residues of the left subunit are numbered with a prime symbol.

### Mutational effects of pore helices on monovalent cation selectivity

The electron density of water molecules in NavAb S178G^NK^, which exhibits the highest Li^+^ selectivity, was observed at the Ser178 position located on the outer edge of the SF. Meanwhile, in the S178A^NK^ with moderately enhanced Li^+^ selectivity, we observed the water molecules besides the Met181 just before the SF vestibule. Accordingly, we hypothesized that the easy influx of fully hydrated ions into the SF increased Li^+^ selectivity. The NavAb Met181 is at the bottom of the extracellular funnel formed by the P2 helix and narrows the pathway to the SF. To increase Li^+^ selectivity by widening the pathway to the SF, the M181A mutations were therefore introduced to NavAb N49K/T206A and its Ser178 mutants (named N49K/T206A-M181A, S178T^NK/TA^-M181A, S178A^NK/TA^-M181A, and S178G^NK/TA^-M181A). Cells expressing S178T^NK/TA^-M181A showed a little current and, rarely, too weak a current of less than 1 nA to evaluate the reversal potential (Figure 7). Additionally, cells exhibiting channel current were scarce in S178A^NK/TA^-M181A and S178G^NK/TA^-M181A, but a credible reversal potential was nonetheless obtained. The resulting *P*_Li_/*P*_Na_ were 0.85 ± 0.03 in N49K/T206A-M181A, 1.09 ± 0.04 in S178A^NK/TA^-M181A and 0.99 ± 0.04 in S178G^NK/TA^-M181A, respectively (Table 2). Li^+^ selectivity increased significantly in N49K/T206A-M181A (*p* < 0.01) and decreased significantly in S178G^NK/TA^-M181A (*p* < 0.01) compared with their pre-mutants. In S178A^NK/TA^-M181A, Li^+^ selectivity was almost unchanged. In other words, the *P*_Li_/*P*_Na_ of those mutants approached 1, indicating non-selectivity for Li^+^ and Na^+^. *P*_Ca_/*P*_Na_ significantly increased in all mutants compared with their pre-mutants. These results indicated that these M181A mutations have caused a loss of monovalent cation selectivity. Met181 of NavAb was thus shown to support the stabilization of the SF and the selection of monovalent and divalent cations. We next focused on the Gln172, toward which the Met181 side chain extends its toe. Then, Gln172 of NavAb N49K/T206A and S178G^NK/TA^ were mutated to evaluate their role in regulating Li^+^ selectivity. The asparagine side chain is shorter than that of glutamine by one methyl group, so the Q172N mutation was expected to weaken or lose the hydrogen bond networks with the Glu177 and the Arg185. Unexpectedly, neither NavAb Q172N mutant conducted any current. This fatal loss of function due to a slight difference in side-chain length suggested a critical role for Gln172 of NavAb in ion permeability and stability of the SF.

**Figure 7.**
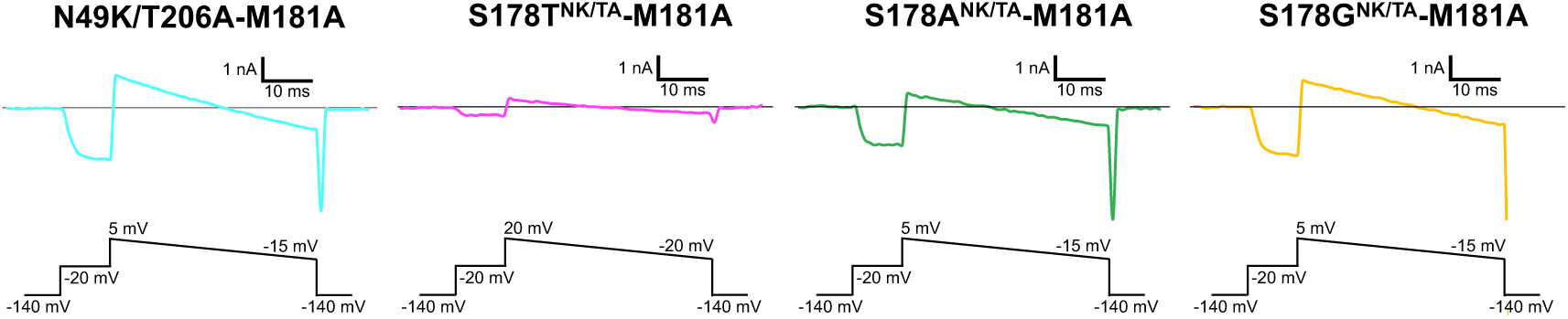
Current traces of NavAb Met181 mutants and Gln172 mutants **A)** Representative current traces of NavAb N49K/T206A-M181A, S178T^NK/TA^-M181A, S178A^NK/TA^-M181A, and S178G^NK/TA^-M181A mutants obtained using the ramp protocol. Currents were measured in 150 mM [Li^+^]_out_ and 150 mM [Na^+^]_in_.

**Table 2.** Relative permeability in NavAb Met181 mutants. All values are indicated as SEM. The p-values represent the results of Student’s t test with the M181A pre-mutants (N49K/T206A, S178^ANK/TA^ and S178^GNK/TA^).

| | $P_{Li}/P_{Na}$ | | $P_K/P_{Na}$ | | $P_{Cs}/P_{Na}$ | | $P_{Ca}/P_{Na}$ | |
| --- | --- | --- | --- | --- | --- | --- | --- | --- |
| N49K/T206A-M181A | 0.84 ± 0.03 | $p < 0.01$ | <0.05* ± 0.03 | NS | <0.01* | $p < 0.01$ | 0.36 ± 0.02 | $p < 0.01$ |
|  | (n = 8) |  | (n = 3) |  | (n = 6) |  | (n = 4) |  |
| S178A<sup>NK/TA</sup>-M181A | 1.09 ± 0.04 | NS | 0.09 ± 0.01 | $p < 0.01$ | <0.05* | NS | 0.41 ± 0.05 | $p < 0.01$ |
|  | (n = 7) |  | (n = 5) |  | (n = 5) |  | (n = 7) |  |
| S178G<sup>NK/TA</sup>-M181A | 0.99 ± 0.04 | $p < 0.01$ | <0.05* ± 0.01 | NS | <0.05* | NS | 0.47 ± 0.05 | $p < 0.01$ |
|  | (n = 3) |  | (n = 3) |  | (n = 3) |  | (n = 4) |  |

\*When the reversal potential was below the deactivation potential, $P_M/P_{Na}$ was noted as < 0.05 which is the value calculated from the lowest measurable reversal potential in each construct.

## Discussion

Various ion selectivity analyses using BacNavs have been reported, but Li^+^ selectivity in all mutants was almost unchanged from that of wild types (*P*_Li_/*P*_Na_ = 0.6∼0.8 in wild types and their mutants) (DeCaen *et al*, 2014; Finol-Urdaneta *et al*, 2014; Naylor *et al*, 2016). In this study, we found that the Ser178 mutations of NavAb enhance Li^+^ selectivity. NavAb S178G^NK/TA^ with the highest Li^+^ selectivity (Li^+^ > Na^+^ > K^+^, Cs^+^) is thought to the first channel that exact belongs to the most Li^+^-selective order of the Eisenman sequence. Following the nomenclature of BacNavs (Yue *et al*, 2002; Tang *et al*, 2014), the NavAb S178G mutants were henceforth collectively referred to as “LivAb” based on their high Li^+^ selectivity (S178G^NK^ and S178G^NK/TA^ were named LivAb^NK^ and LivAb^NK/TA^, respectively).

Hydration water exchanges are crucial for ion permeation in Navs (Ahern *et al*, 2016). The diameter of the SF of Navs is sufficiently wide that two to three Na^+^ ions are accommodated in the SF and permeate in a hydrated or partially dehydrated state (Chakrabarti *et al*, 2013; McCusker *et al*, 2012; Payandeh *et al*, 2011; Shaya *et al*, 2014; Tsai *et al*, 2013; Ulmschneider *et al*, 2013). Referring to the crystal structure of NavAb, the oxygen atoms in the main and side chains in the SF form four hydration water exchange sites (Figure 1A) (Payandeh *et al*, 2011). When fully hydrated Na^+^ in bulk solution approaches the outer edge of the SF in BacNavs, it first interacts with protein via water molecules (Naylor *et al*, 2016). In the case of NavAb, entering into the SF requires the Na^+^ to release two hydration waters, which are subsequently replaced by the oxygen atoms of the side chains of Ser178 and Glu177 (Corry & Thomas, 2012). Na^+^ passes through the following hydration water exchange sites formed by the main-chain carbonyls of Leu176 and Thr175 and then enters the central cavity (Payandeh *et al*, 2011; Chakrabarti *et al*, 2013). That is, changes in the number and location of hydration water exchanges in the SF will significantly affect the permeation efficiency of ions with different properties.

Hydration water exchange is one of the rate-limiting steps of ion transfer. Na^+^ and Li^+^ are monovalent and single-atom cations, and the ionic radius of Li^+^ (0.60 Å) is smaller than that of Na^+^ (0.95Å). Due to its small radius, Li^+^ has a stronger electrostatic interaction with the negative dipole of water molecules than Na^+^, resulting in a slower rate of hydration water exchange(Hille, 2001). Hence, the more hydration water exchange sites formed by hydroxyl groups in the SF, the slower the rate of Li^+^ permeation with a strong interaction force and the lower its permeability. In this study, the hydroxyl groups that form the first hydration water exchange site in the SF were lost in NavAb S178A^NK/TA^ and S178G^NK/TA^ (LivAb^NK/TA^) compared with N49K/T206A (Figure 8). This decrease in hydration water exchanges in the SF may have worked in advantage of the permeation of Li^+^, which has slower exchange rate.

**Figure 8.**
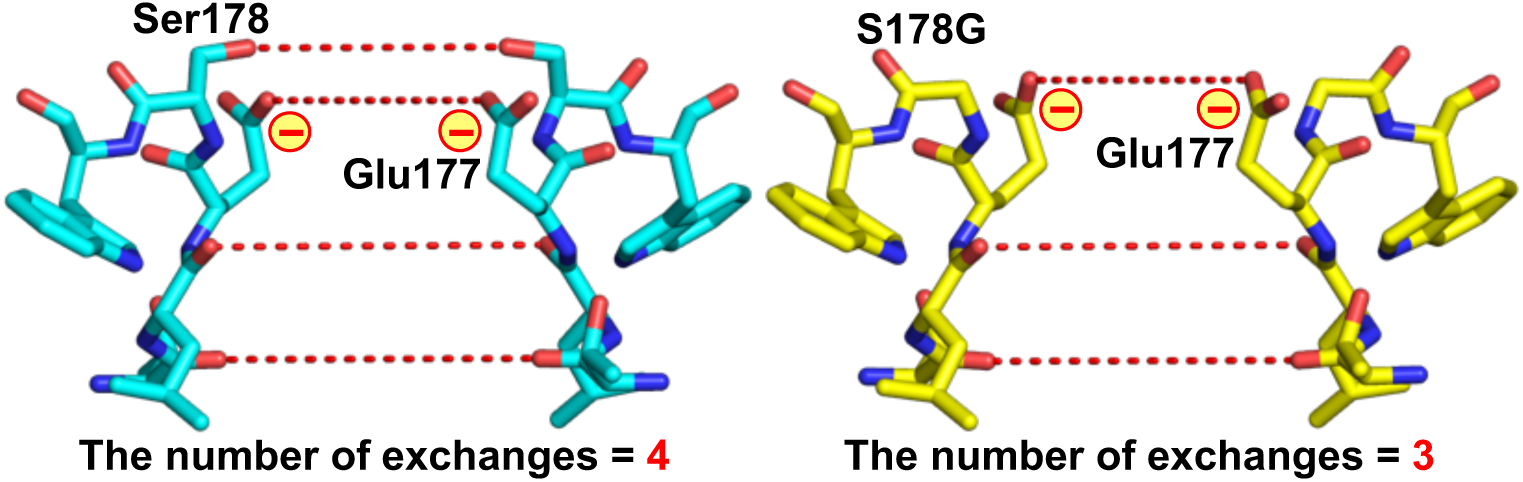
The selectivity filter of NavAb N49K and S178G^NK/TA^ (LivAb^NK/TA^) mutants Side view of the residues 175-179^th^ constituting the SF of N49K (left, PDB code 8H9W) and S178G^NK/TA^ (LivAb^NK/TA^) (right, PDB code 9UC3). The red dashed lines represent the hydration water exchange sites which ions pass through, formed by the oxygen atoms of the main and side chains of the SF.

The reduction of hydration water exchanges in the SF alone does not increase the permeability of Li^+^ to exceed that of Na^+^ because it simultaneously increases the permeability of both Na^+^ and Li^+^. According to the Eisenman sequence, which divides the monovalent cation selectivity in 11 ways, the selectivity of Li^+^ only exceeds that of Na^+^ when the electrostatic force interacting with ions is most potent (the order Ⅺ: Li^+^ > Na^+^ > K^+^ > Cs^+^) (Hille, 2001; Eisenman, 1962). The most potent electrostatic interaction site of NavAb is comprised of the side chain of Glu177, which is only negatively charged in the SF (TLESW) (Payandeh *et al*, 2011). Glu177 of NavAb and corresponding residues of eukaryotic Navs and Cavs are known to bind cations as the most strongly electrostatic site and facilitate selective permeation of ions (Boiteux *et al*, 2014; Chakrabarti *et al*, 2013; Favre *et al*, 1996; Ke *et al*, 2014; Lipkind & Fozzard, 2000; Xia *et al*, 2013). In the NavAb S178A and S178G (LivAb) mutants, the loss of the hydration exchange site formed by the side chain of Ser178 would expose Glu177 directly to the bulk solution. The strength of ionic bond depends on the electrostatic force and, inversely, on the sum of the radii of two particles. The exposed electrostatic site is, therefore, advantageous for the ionic bond with Li^+^, which has a smaller ionic radius and a stronger positive charge than Na^+^. The fact that the exposed Glu177 side chain enhanced Li^+^ selectivity corroborates the correctness of Eisenman sequence. In addition, in the NavAb S178G (LivAb), with the glycine mutation widening the SF vestibule, ions from bulk solution were thought to easily approach the exposed electrostatic site directly, leading to a reversal of Li^+^ and Na^+^ permeability. In the crystal structures of S178A^NK^ and S178G^NK^ (LivAb^NK^), the electron density of water molecules were observed at the SF vestibule. It explains the high accessibility of hydrated ions in bulk solution to the most electrostatic sites in these mutants.

Here, we should focus on why the NavAb S178G^NK/TA^ mutant retains low Ca^2+^ selectivity, with *P*_Ca_/*P*_Na_ < 0.01, while enhancing Li^+^ selectivity, because Ca^2+^ has strong electrostatic force than monovalent cations. We previously reported the only identified prokaryotic Cav, CavMr (Irie, 2021; Shimomura *et al*, 2020). CavMr functions as a homotetramer, similar to prokaryotic Navs, and the SF sequence of CavMr is “TLEGW”, the same as that of S178G^NK/TA^ (LivAb^NK/TA^). The glycine residue in the SF (TLEGW) of CavMr plays a key role in Ca^2+^ selectivity, and a single-point mutation of its glycine to serine decreases *P*_Ca_/*P*_Na_ about 18-fold. In contrast, this study found that S178G mutation of NavAb affected only Li^+^ selectivity, not Ca^2+^ selectivity. However, all the NavAb M181A mutants were converted into non-selective channels with moderate Li^+^ permeability, equivalent to Na^+^, and approximately 10-fold Ca^2+^ selectivity. This result suggests that Met181 of NavAb is important in sorting monovalent and divalent cations (Li^+^ and Na^+^ v.s. Ca^2+^ in this case).

The corresponding position of NavAb Met181 in CavMr is aspartate (Shimomura *et al*, 2020), and the same is true for artificial Cavs (CachBac and CavAb) (Yue *et al*, 2002; Tang *et al*, 2014). Their aspartate side chains are considered to affect cation permeability through electrostatic interactions. On the other hand, if only the electrostatic repulsion is required for cation selectivity by Met181 in NavAb, Li^+^ should also be excluded by its stronger positive charge than Na^+^, as should Ca^2+^. In other words, factors other than electrostatic interaction are involved in the monovalent cation selectivity by NavAb Met181. Around the SF, extensive interactions are formed within the same subunit or between neighboring subunits (Payandeh *et al*, 2011). As for the crystal structure of NavAb, it has been known that Met181 and Gln172, located at the pore helices, also form part of the network; however, their roles have only been rarely experimentally evaluated. In this study, the small number of cells exhibited activity and stable currents in the NavAb M181A mutants, particularly in S178T^NK/TA^-M181A. It proposes the idea that Met181 of NavAb promotes monovalent cation selectivity through SF stabilization, and that this effect is more substantial in the S178T mutant. This combination effect of Met181 and S178T may be a reason for the increase in Li^+^ selectivity in S178T^NK/TA^, which retain the first hydration water exchange site. To better understand the molecular basis of ion selectivity of Navs, detailed analyses of these extensive interactions are required, especially those between the SF and pore helices.

In summary, the present study elucidated the mechanisms of high Li^+^ selectivity in Navs using a BacNav. The strong electrostatic force in the SF mainly contributes to the high Li^+^ selectivity, and the smaller number of hydration exchanges promotes Li^+^ permeation in Navs. Differences in ionic properties, such as ionic radius, electrostatic energy, and hydration water exchange rate, were related to the Li^+^ selectivity of Navs. It was also revealed that the SF stabilization by the extracellular funnel and pore helices supports the selective permeation of monovalent cations. These discoveries of the molecular basis of the high Li^+^ selectivity of Navs and the creation of LivAb, a novel “Li channel”, provide direction for drug development targeting various neurological disorders and become the cue to understanding the biological effects of lithium.

## Methods

### Site-directed mutagenesis and construction of NavAb mutants

DNA constructs were produced as previously reported (Irie *et al*, 2023). The NavAb mutated DNAs were subcloned into the modified pBiEX-1 vector (Novagen; 71234-3CN) that was modified by replacing the fragment from NcoI site (CCATGG) to SalI site (GTCGAC) in multicloning site with the sequence “CCATGGGCAGCAGCCATCATCATCATCATCACAGCAGCGGCCTGGTGCCG CGCGGCAGCCATATGCTCGAGCTGGTGCCGCGCGGCAGCGGATCCTAAGT CGAC” (Irie *et al*, 2018). The NavAb mutated DNAs were subcloned between BamHI and SalI sites to add the N-terminal His tag and thrombin cleavage site. The polymerase chain reaction accomplished site-directed mutagenesis using PrimeSTAR® Max DNA Polymerase (Takara Bio). All clones were confirmed by DNA sequencing.

### Electrophysiological measurement in insect cells

The recordings were performed using SF-9 insect cells (ATCC catalogue number CRL-1711), which were grown in Sf-900™ II medium (Gibco) supplemented with 1% 100× Antibiotic-Antimycotic (Fujifilm-Wako) at 27°C. Cells were transfected with target channel-cloned pBiEX-1 vectors and enhanced green fluorescent protein (EGFP)-cloned pBiEX-1 vectors using polyethyleneimine (PEI) reagent (Cosmo bio). PEI was solubilized in distilled water, and 1 mg/mL PEI solution pH was adjusted to 7.0 with NaOH. First, the channel-cloned vector (3.3 μg) was mixed with 1.65 μg of the EGFP-cloned vector in 150 μL of the culture medium. Next, 15 μL of a 1 mg/mL PEI solution was added, and the mixture was incubated for 10 min before the transfection mixture was gently dropped onto the cultured cells. After 24-48 h incubation, the cells were used for electrophysiological measurements.

For reversal potential measurements to determine the relative permeabilities of Na^+^ and other cations, the internal pipette solution contained 35 mM NaCl, 115 mM NaF, 10 mM EGTA, and 10 mM HEPES-NaOH (pH 7.4). The pipette solution was mixed with the acidic solution (35 mM NaCl, 115 mM NaF, 10 mM EGTA, and 10 mM HEPES) and the basic solution (35 mM NaOH, 115 mM NaF, 10 mM EGTA, and 10 mM HEPES) to adjust the pH to 7.4. For the evaluation of cation selectivity, the Li^+^ extracellular solution was mixed with the acidic solution (150 mM LiCl, 10 mM HEPES, 10 mM glucose, 2 mM CaCl_2_ ) and the basic solution (150 mM LiOH, 10 mM HEPES, 10 mM glucose, 2 mM CaCl_2_) to adjust pH 7.4. The K^+^, Cs^+,^ and Ca^2+^ extracellular solutions were similar, except that all the LiCl and LiOH were replaced by an osmotically equivalent amount of these cation salts and basic components. Cancellation of the capacitance transients and leak subtraction were performed using a programmed P/10 protocol delivered at -140 mV. The bath solution was changed using the Dynaflow^®^ Resolve system. All experiments were conducted at 25 ± 1°C using a whole-cell patch clamp mode with a HEKA EPC 10 amplifier and Patch master data acquisition software (v2×73). Data export was performed using Igor Pro 9.05 and NeuroMatic (v3.0s) (Rothman & Silver, 2018). All sample numbers represent the number of individual cells used for each measurement. Cells with a smaller leak current than 1 nA were used for data collection. When any outliers were encountered, they were excluded if any abnormalities were found in other measurement environments, and they were included if no abnormalities were found. All results are presented as mean ± standard error. The graph data were plotted using Microsoft Excel (v16.97).

### Calculation of ion selectivity by the GHK equation

The intracellular and extracellular solutions were arbitrarily set to determine the ion selectivity of each channel. The reversal potential at each concentration was measured by applying a ramp pulse to the membrane potential. A 10-ms depolarization stimulus was inserted for each measurement to confirm the cell’s condition. The permeation ratio of Ca^2+^ and Na^+^ (*P*_Ca_/*P*_Na_) was calculated by substituting the obtained reversal potential (*E*_rev_) into the expression derived from the GHK equation (Frazier, George, and Jones, 2000):

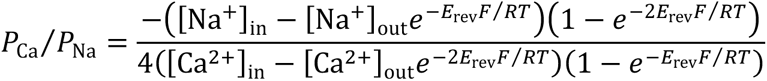

Where R, T, F, and *E*_rev_ are the gas constant, absolute temperature, Faraday constant, and reversal potential, respectively.

The permeation ratio of monovalent cations (*P*_M_/*P*_Na_, M represents Li^+^, K^+^, or Cs^+^) was calculated by substituting the obtained reversal potential and *P*_Ca_/*P*_Na_ into the expression derived from the GHK equation (Lopin *et al*, 2012):

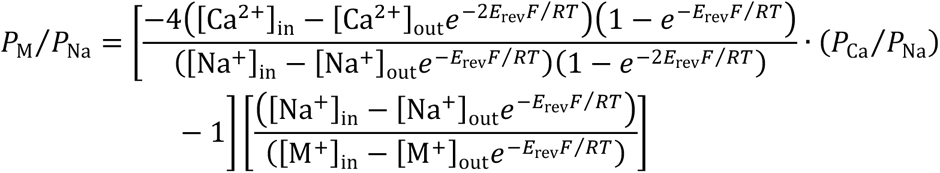

### Protein expression and purification

Proteins were expressed in the *Escherichia coli* KRX strain (Promega). Cells were grown at 37°C to an OD_600_ of 0.6, induced with 0.1% rhamnose (Fujifilm-Wako), and grown for 16 h at 20°C. The cells were suspended in TBS buffer (20 mM Tris-HCl pH 8.0, 150 mM NaCl) and lysed using LAB1000 (SMT Co., LTD.) at 12,000 psi. Low-speed centrifugation removed cell debris (12,000×g, 30 min, 4°C). Membranes were collected by centrifugation (100,000×g, 1 h, 4°C) and solubilized by homogenization in TBS buffer containing 30 mM n-dodecyl-β-D-maltoside (DDM, Anatrace). After centrifugation (40,000×g, 30 min, 4°C), the supernatant was loaded onto a HIS-Select^®^ Cobalt Affinity Gel column (Sigma). The protein bound to the cobalt affinity column was washed with 10 mM imidazole in TBS buffer containing 0.05% lauryl maltose neopentyl glycol (LMNG, Anatrace) instead of DDM. After washing, the protein was eluted with 300 mM imidazole, and the His tag was removed by thrombin digestion (overnight, 4°C). Eluted protein was purified on a Superdex-200 column (Cytiva) in TBS buffer containing 0.05% LMNG.

### Crystallization and structural determination

Before crystallization, the purified protein was concentrated to ∼10mg ml^−1^ and reconstituted into a bicelle solution (Faham *et al*, 2005), containing a 10% bicelle mixture at 2.8:1 (1,2-dimyristoyl-sn-glycero-3-phosphorylcholine [DMPC, Anatrace]: 3-[(3-cholamidopropyl) dimethylammonio]-2-hydroxypropanesulfonate [CHAPSO, DOJINDO]). The NavAb 10% bicelle was mixed in a 20:1 ratio. Prepared proteins were crystallized by sitting-drop vapor diffusion at 20°C by mixing 300-nl volumes of the protein solution (8–10 mg/ml) and the reservoir solution (9%–11% polyethylene glycol monomethyl ether [PEG MME] 2000, 100 mM sodium chloride, 100 mM magnesium nitrate, 25mM cadmium nitrate and 100 mM Tris-HCl, pH 8.4) with mosquito LCP (STP Labtech). The crystals were grown for 1 to 3 weeks, and after growing, the crystals were transferred into the reservoir solution without cadmium nitrate, which was replaced with the following cryoprotectant solutions. The cryoprotectant solution contains 11% PEG MME 2000, 100 mM Tris-HCl pH 8.4, 2.5 M sodium chloride, 100 mM calcium nitrate, and 20% (v/v) DMSO.

All data were automatically collected at BL41XU and BL45XU of SPring-8 and merged using the ZOO system (Hirata *et al*, 2019). Data were processed using the KAMO system (Yamashita *et al*, 2018) with XDS (version 20220110) (Kabsch, 2010). The datasets of NavAb S178T mutants (S178T^NK^ and S178T^NK/TA^) were obtained from a single crystal, while the data sets of NavAb S178A mutants (S178A^NK^ and S178A^NK/TA^) and S178G^NK^ were obtained from four and three crystals, respectively. Analyses of the data with the STARANISO server (Global Phasing Ltd) revealed severely anisotropic crystals. The data sets were, therefore, ellipsoidally truncated and rescaled to minimize the inclusion of poor diffraction data.

A molecular replacement method using PHASER (McCoy *et al*, 2007) provided the initial phase using the structure of the NavAb N49K mutant (pdb code: 8H9W) as the initial model. The final model was constructed in COOT (version 0.9.2) (Emsley *et al*, 2010) and refined in REFMAC5 (Murshudov *et al*, 1997) and Phenix (version 1.18) (Adams *et al*, 2010). The CCP4 package (version 7.0.078) (Winn *et al*, 2011) was used for structural analysis. Data collection and refinement statistics for all crystals are summarized in Supplementary Table 1. All figures were prepared using the program PyMOL 2.5.4 (Schrödinger, LLC, 2015).

### Statistics

Electrophysiological data were analyzed using Microsoft Excel (v16.97). All results are presented as the mean with standard error (SEM). Student’s t-test was used to estimate statistical significance between the NavAb M181A mutants and their pre-mutants, and *p* < 0.05 was considered statistically significant.

## Data availability

The data that support this study are available from the corresponding authors upon request. The structural data generated in this study have been deposited in the Protein Data Bank under accession codes 8H9W (NavAb N49K mutant in calcium), 9UC1 (NavAb S178T^NK^ mutant), 9UC2 (NavAb S178A^NK^ mutant), 9UC3 (NavAb S178G^NK^ mutant [LivAb^NK^]), and 9UC4 (NavAb S178T^NK/TA^ mutant). The structure of the NavAb N49K mutant for the initial model of molecular replacement was available in the Protein Data Bank under accession code 8H9W.

## Author contributions

Yuki K. Maeda: Conceptualization; Resources; Data curation; Formal analysis;

Validation; Investigation; Writing - original draft, Writing - review and editing

Kentaro Kojima: Data curation

Tomoe Y. Nakamura-Nishitani: Writing - review and editing

Toru Nakatsu: Funding acquisition; Writing - review and editing

Katsumasa Irie: Conceptualization; Funding acquisition; Resources; Data curation;

Software; Formal analysis; Supervision; Validation; Investigation; Visualization; Writing

- original draft; Project administration; Writing - review and editing

## Acknowledgements

The synchrotron radiation experiments were performed at BL41XU and BL32XU in SPring-8 with the approval of the Japan Synchrotron Radiation Research Institute (JASRI) (Proposal numbers 2016B2721, 2017B2735, and 2018B2710). We thank the beamline staff for their excellent facilities and support. This research was partially supported by Platform Project for Supporting Drug Discovery and Life Science Research (Basis for Supporting Innovative Drug Discovery and Life Science Research [BINDS]) from AMED under Grant Numbers JP22ama121001 and JP23ama121001. This work was supported by Grants-in-Aid for Scientific Research (17K17795, 20K09193, and 24K02168) and SEI Group CSR Foundation, Takeda Science Foundation, Institute for Fermentation, and the Salt Science Research Foundation.

---

**Supplementary Table 1.**
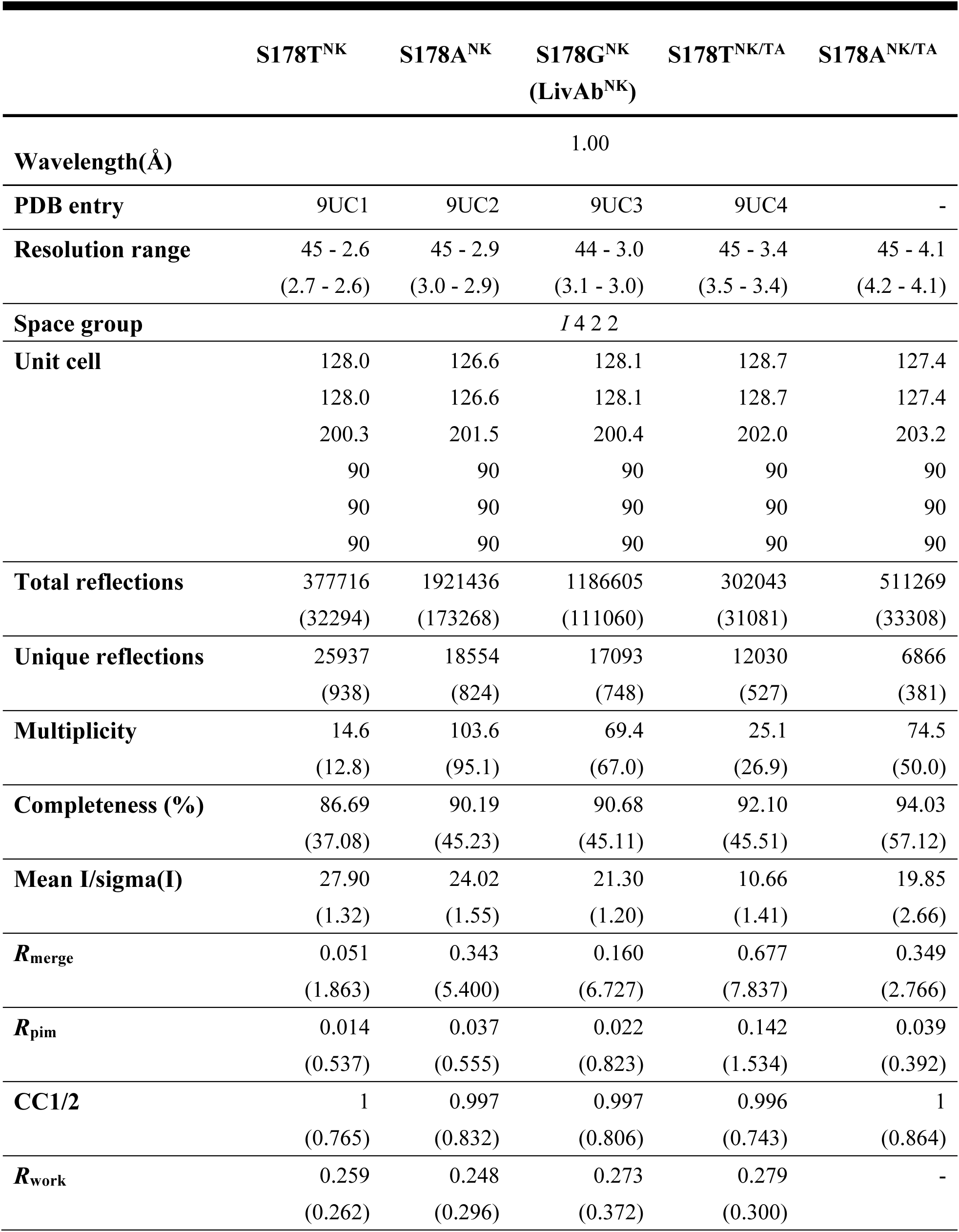
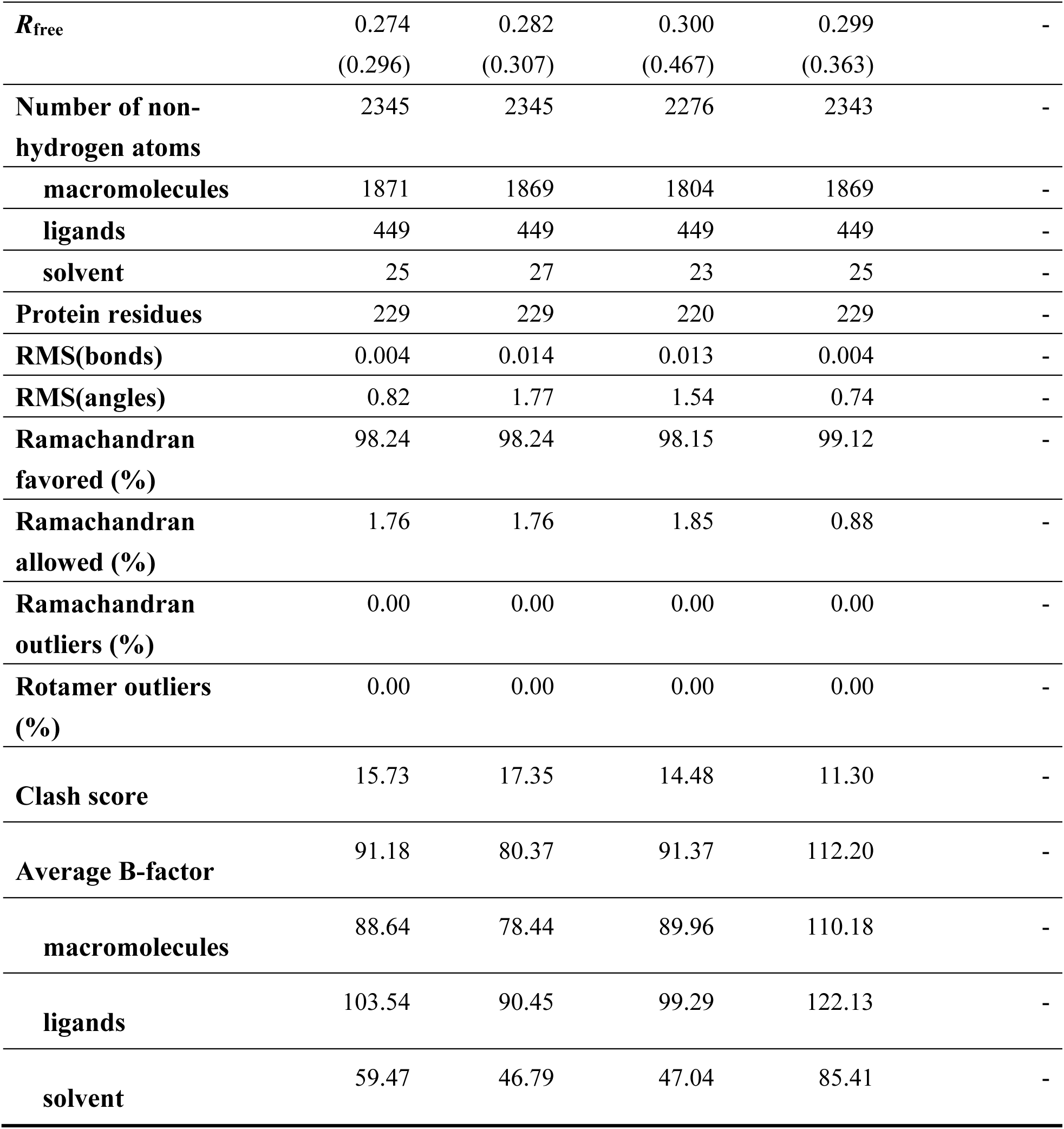
Data collection and refinement statistics.

